# *In vitro* analysis of the effects of acetaminophen, ibuprofen and aspirin on motility of female adult *Onchocerca volvulus* worms

**DOI:** 10.1101/2021.10.20.465198

**Authors:** Kenneth B. Otabil, Rhoda K. Antwi, Prince Nyarko, Theophilus N. Babae, Blessing Ankrah, Emmanuel J. Bart-Plange, Rexford Kyei, Joseph Ameyaw, Joseph G. Bamfo, Samuel F. Gyasi, Henk DFH Schallig

## Abstract

In view of the very ambitious global timelines for elimination of onchocerciasis in 2030, the search for alternative antifilarials cannot depend on drug development from scratch, and repurposed drugs offer cheaper and faster alternatives. Previous studies had demonstrated the presence and potential expression of amidase, an enzyme that can be targeted by repurposed analgesics, *in Onchocerca volvulus worms*. The aim of this study was to determine the effects of acetaminophen, ibuprofen and aspirin on the motility of adult *O. volvulus* worms. In total, thirty (30) female *O. volvulus* worms were exposed to acetaminophen, ibuprofen and aspirin in concentrations of either 5mg/ml, 2.5mg/ml, 1.25mg/ml or 0.63mg/ml control in duplicates. Worm motility was observed and recorded using the WormAssay software and a darkfield imaging apparatus starting on day 2 of incubation and ending on day 8. Acetaminophen, ibuprofen and aspirin inhibited *O. volvulus* motility by 2-fold compared to control in the first 24 hours of drug exposure. However, an extended exposure of the worms to these test drugs rather improved the motility of the worms. The study has demonstrated that a 24-hour exposure of *O. volvulus* worms to the analgesic drugs studied, results in significant inhibition in worm motility compared with the control group, but extended duration of exposure led to an enhancement in motility of the previously immobile worms. This finding supports the idea that aspirin and ibuprofen may have some longevity enhancing properties. Further research on the utility of these analgesics as possible anti-filarial drugs is thus warranted.

## Introduction

Onchocerciasis (river blindness) is a neglected tropical disease (NTD), caused by the parasite *Onchocerca volvulus* and transmitted by blackflies [1]. Currently, the disease is still a major public health concern even after years of both vector control and Mass Drug Administration (MDA) with ivermectin [2]. It is a chronic cutaneous and ocular disease characterized by skin nodules, severe itching and ocular lesions that can progress to partial or total blindness [3]. Globally, about 21 million people are infected and a further 200 million people at risk of contracting the disease according to the 2017 Global Burden of Disease Study [4]. Sub-Saharan Africa (SSA) bears approximately 99% of the disease burden [5]. In Ghana, recent updates demonstrated that 15 of its 16 regions (120 districts) are endemic for onchocerciasis, with more than 7 million people at risk of contracting the disease [6].

The key strategy for onchocerciasis control has been the use of ivermectin for the past 10 years as an effective microfilaricidal drug with no significant macrofilaricidal effects. However, some worrying reports from studies in West Africa have provided evidence of the development of sub-optimal response of *O. volvulus* parasites to ivermectin [7]–[9]. Aside this challenge, as ivermectin is purely microfilaricidal and does not kill adult worms [10], control programmes have had to administer the drug for many years in order to interrupt the transmission of the disease in endemic communities [11]. A major breakthrough in anti-filarial therapy was the discovery of anti-*Wolbachia* antibiotics such as doxycycline, which demonstrably had adulticidal effects [12]. However, logistical challenges associated with mass administration of antibiotics prevents the deployment of these antibiotics at the community level (Hoerauf, 2008; Hugues, Nana-Djeunga et al., 2014; Debrah et al., 2015). As a result of these difficulties, coupled with the suspension of the World Health Organization (WHO) sponsored Onchocerciasis Control Programmes in endemic communities since 2002, there seems to be a resurgence of the disease in some tropical foci [14].

It is against this background that the global health community, led by the WHO through a recently published NTD roadmap for 2021–2030 [15] has set elimination of transmission (EOT) targets for onchocerciasis, with 12 countries verified for EOT by 2030. Nevertheless, in some highly endemic foci, it might be impossible to achieve elimination of the disease with a sole reliance on ivermectin [16]. Therefore, the search for a microfilaricide and alternative drugs against onchocerciasis remains a top research priority [17].

The discovery and development of new drugs requires a lot of time (10–15 years), complex and expensive processes, and the success rate has been low, only 2.01% [18]. This makes it less advantageous for current onchocerciasis control efforts to depend on development of new drugs from scratch in view of the ambitious EOT timelines of 2030. In this regard, drug repurposing may be the best strategy to discover effective adulticidal drugs from existing approved drugs since these have already passed preclinical and clinical stages [19], [20] and this stands out as an attractive option to find alternative antifilarial options for onchocerciasis.

A major discovery was made by Cotton *et al*., [21] when they sequenced the entire genome of *O. volvulus* parasites and its *Wolbachia* endosymbiont. They provided a list of gene families that had key functions, which could potentially be exploited as targets for repurposed drugs against the parasite [21]. The investigators also identified enzymes (and their gene targets) that are potentially chokepoint enzymes and may thus be essential for *O. volvulus* survival [21]. One of the top 16 *O. volvulus* gene targets identified was the OVOC4110 gene which potentially codes for amidase, which may be critical in metabolic pathways of *O. volvulus* [21]. The authors opined that analgesics such as acetaminophen may block the action of amidase and consequently disrupt critical metabolic pathways in the worm [21]. As yet, no study has confirmed the expression of amidase in *O. volvulus* worms or the effects of these analgesics on adult *O. volvulus* worms, to the best of our knowledge. For the present study, we investigated the effects of acetaminophen, ibuprofen and aspirin on adult *O. volvulus* worms with the goal of assessing the potential of these analgesics as repurposed, macrofilaricidal drugs to aid in the control of onchocerciasis.

Acetaminophen (paracetamol), an analgesic and antipyretic, is the most widely used over-the-counter medication in the most countries [22]. The mechanism of action of acetaminophen is complex and includes the effects of both the peripheral (Cyclo-oxygenase inhibition), and central (Cyclo-oxygenase, serotonergic descending neuronal pathway, L-arginine/Nitric Oxide pathway, cannabinoid system) antinociception processes and Redox mechanisms [22]. Though, some studies have reported some antimicrobial effects of acetaminophen (AL-Janabi, 2010), the direct activity of acetaminophen against helminths in general and *O. volvulus* in particular has not been demonstrated by any study till date, to the best of our knowledge. Cotton et al. (2016) however speculated that acetaminophen might bind to amidases in *O. volvulus* and potentially cause the death of the worms, hence its choice in the present study.

Ibuprofen is also a medication in the nonsteroidal anti-inflammatory drug (NSAID) class that is used for treating pain, fever, and inflammation [24]. Like other NSAIDs, it works by inhibiting the production of prostaglandins by decreasing the activity of the enzyme cyclooxygenase [25] though it might be a weaker anti-inflammatory drug than other NSAIDs [25]. The drug was chosen for the present study because aside being an analgesic, some antimicrobial activity has been reported by previous authors [23], [26]. However, till date no study has fully investigated the direct effects of ibuprofen on *O. volvulus* survival.

Aspirin (acetylsalicylic acid, ASA), an analgesic, antipyretic and anti-inflammatory drug, has emerged as an interesting option for repurposing drugs against parasites [18]. ASA may also be used for secondary prevention of stroke and acute cardiac events. ASA acts on the two cyclooxygenase isoforms (Cyclo-oxygenase 1and 2) via acetylation of serine 532 in the active site; inhibits the action of both enzymes and prevents the formation of prostaglandins from arachidonic acid [27]. Epidemiological and clinical studies has shown that ASA also reduces the incidence of epithelial tumours by the acetylation of multiple proteins including transcription factors, cytoskeleton proteins, stress response proteins (including the heat shock proteins, HSPs), membrane proteins, among others [28], [29], suggesting other mechanisms and molecular targets. Some studies have evidenced antiparasitic activity, for instance, Singh and Rathaur [30] demonstrated that ASA improved the antifilarial activity of diethylcarbamazine [30]. Aspirin also demonstrably affects dynamics of the actin cytoskeleton and decreases amebic movement in *Entamoeba histolytica (López-Contreras et al., 2013)*.

The aim of the study therefore was to determine the effects of acetaminophen, ibuprofen and aspirin on the survivability of adult *O. volvulus* worms. This was important because there is the need to quickly discover and develop effective macrofilaricidal alternatives to complement ivermectin if global elimination targets of onchocerciasis are to be achieved.

## Materials and Methods

### Patient recruitment and nodulectomies

*Onchocerca* nodules were obtained from 11 participants who had consented to participate in this study by signing/thumbprinting a written, informed consent form. The study was approved by the Institutional Ethics Committee of the Kintampo Health Research Centre in Ghana (Approval number: KHRCIEC/2018-18). The participants were recruited from 7 onchocerciasis endemic communities during a cross-sectional, epidemiological field survey in November 2020 in the Bono Region of Ghana. A total of 560 individuals from all 7 communities were screened for cutaneous nodules (onchocercoma) by palpation [32] by the medical team and eligible participants with palpable nodules, with no condition requiring long-term medication, were recruited and scheduled for nodulectomies. Using local anaesthesia, palpable nodules were surgically removed from the patients by the medical team. Participants were then transported to their homes and wound care provided at the nearest health care facility until the wound was healed under the supervision of the medical team. The excised nodules were preserved in 0.85% normal saline in 20 ml plastic vials, placed on ice, transported to the laboratory and stored at room temperature on the laboratory bench.

### Collagenase digestion of excised nodule

In order to isolate the adult worms contained in the nodules, the latter were digested using the collagenase technique [33]. Briefly, each nodule was incubated for 48 hours at 37°C (depending on the nodule’s weight) in 5ml of Gibco’s high glucose Dulbecco’s Modified Eagle’s Medium (DMEM, Gibco, 12100061) containing type I collagenase (Sigma Aldrich, SCR103) at a final concentration of 2.25 mg/ml. The product of digestion (the worm mass and digested human tissues constituting the nodule) was placed in a Petri dish containing 15 ml of DMEM, sodium bicarbonate (Dainess, Ghana), and supplemented with Penicillin-Streptomycin (Gibco, 15140122) at a final concentration of 2 mg/ml. Individual worms were isolated under a dissecting microscope using entomological forceps and needles. Each entire and living worm (dead or calcified and incomplete or broken worms were counted but discarded from the further process) was then examined for sex. Entire male worms were individually frozen for further studies. Thirty (30) female worms were examined for activity in order to include them in the live worm assays after 24 hours of separation from nodules. Males were excluded from this study because initial culture of the male worms indicated an unusually fast decay in DMEM.

### Drug preparation

Stock solutions of acetaminophen, ibuprofen, aspirin were prepared by grinding 500 mg of the commercially available drugs [Paracetamol (M&G Pharmaceuticals, Ghana); Ibuprofen (Letap Pharmaceuticals, Ghana); Aspirin (M & A Pharmaceuticals, UK)] in a breakable crucible and dissolved in 50 ml of DMEM solution in a 250 ml beaker. Subsequently, concentrations of 5 mg/ml, 2.5 mg/ml, 1.25 mg/ml and 0.63 mg/ml were prepared using two-fold serial dilution following a protocol developed by Al-Janabi [23]. The concentration of acetaminophen, ibuprofen and aspirin in micromolars (mM) were; Acetaminophen (5 mg/ml=33.08mM, 2.5 mg/ml=16.54 mM, 1.25 mg/ml = 8.27 mM and 0.63 mg/ml = 4.17 mM), Ibuprofen (5 mg/ml = 24.24 mM, 2.5 mg/ml = 12.12 mM, 1.25 mg/ml = 6.06 mM and 0.63 mg/ml = 3.06 mM), Aspirin (5 mg/ml = 27.75 mM, 2.5 mg/ml = 13.88 mM, 1.25 mg/ml = 6.94 mM and 0.63 mg/ml = 3.50 mM). The drug dilutions were stored at 4°C in a refrigerator for 24 hours prior to usage in the O. volvulus assays.

### O. volvulus worm culture nodule

The worms were cultured following established protocols [34] with a few modifications. The experiment was set up by adding a single female adult *O. volvulus* worm to each well of 6-well plates containing 5 ml of DMEM +10% fetal bovine serum (Gibco, A3840002) + 100 u/ml penicillin + 100 ug/ml streptomycin + sodium bicarbonate (2 g/1). The worms were then incubated at 37 °c in a gas phase of 5 % CO_2_ in air for 24 hours prior to the addition of the analgesics to adapt them to the culture medium. After 24 hours of incubation in the above medium, 1 ml of each concentration of analgesics (5, 2.5, 1.25 and 0.63 mg/ml) was pipetted into a previously appropriately labelled wells containing the pre-incubated *O. volvulus* worms in the medium. Two (2) wells with DMEM but no analgesics added were used as the control for this study. The worms were again incubated at 37 °C in a gas phase of 5% CO_2_ in air for 8 days with motility reading performed daily starting after 24 hours. Close examination of worms was performed to identify signs of disintegration and the medium changed every other day as colour changes indicated depletion of glucose in the culture medium.

### The WormAssay and the darkfield imaging apparatus

The WormAssay software previously developed by Marcellino et al. (2012) for antifilarial screening of *Brugia malayi* was repurposed in the present study for *O. volvulus* [36]. The WormAssay is an inexpensive system for quantifying parasite movement based on worm motility. The apparatus uses a commodity video camera, computer and a newly developed free and open source software application [35] to provide quantitative measurements of parasite motility on entire plates. The application is designed to process multiple wells simultaneously without user interaction and this automatically identifies each well in the plate and labels the output data accordingly. This system can be used to assay large parasites such as filarial nematodes as well as other macroparasites. WormAssay’s automation of the video capture step and lack of need for any interaction with the computer software during scoring differentiates it from other existing motion-based schemes. For the video capture system, a dark field macroscopic imaging apparatus based on the design of Marcellino et al. (2012) was developed with slight modifications.

### Measurement of motility using the WormAssay software and darkfield imaging system

Before the addition of the test drugs, the motility of the female worms to be used were observed by competent laboratory technicians (parasitologists) and judged to be of sufficient motility to include in the assays. Using the WormAssay and the capture device, each 6-well microplate was observed and scored for motility. One (1) minute video recordings using the Lucas-Kande Optical Flow algorithm were taken of each plate and mean optical flow movement units for each worm were converted to percent motility by normalization in Graphpad Prism. The effective concentration to reduce mean motility levels to 50% (EC50) of control values were also determined using Graphpad Prism. The full profile of % mean motility scores (MMS) were illustrated graphically and statistical comparisons made.

### Statistical analyses

Statistical analysis was performed using Graphpad Prism (version 8.2.1). In plotting the dose response curves, all movement units were normalized and the EC50 (the concentration of a drug that gives half-maximal response was calculated for acetaminophen, ibuprofen and aspirin) were determined. For the graphical plots, the mean of the normalized movement units was plotted against the concentration or the day depending on the type of plot. Differences in mean motility scores between groups were calculated using the Mann-Whitney test with differences considered to be statistically significant at the 95 % confidence interval if P < 0.05. The standard deviations (SD) of the mean motility scores were determined and subsequently converted to % SD for consistency.

## Results

The results of the effects of acetaminophen, ibuprofen and aspirin on the survival of adult female *O. volvulus* worms are presented in Figures 1–5. The indicator of survival of *O. volvulus* female worms in this study was the percentage mean motility scores (% MMS).

**Figure 1:**
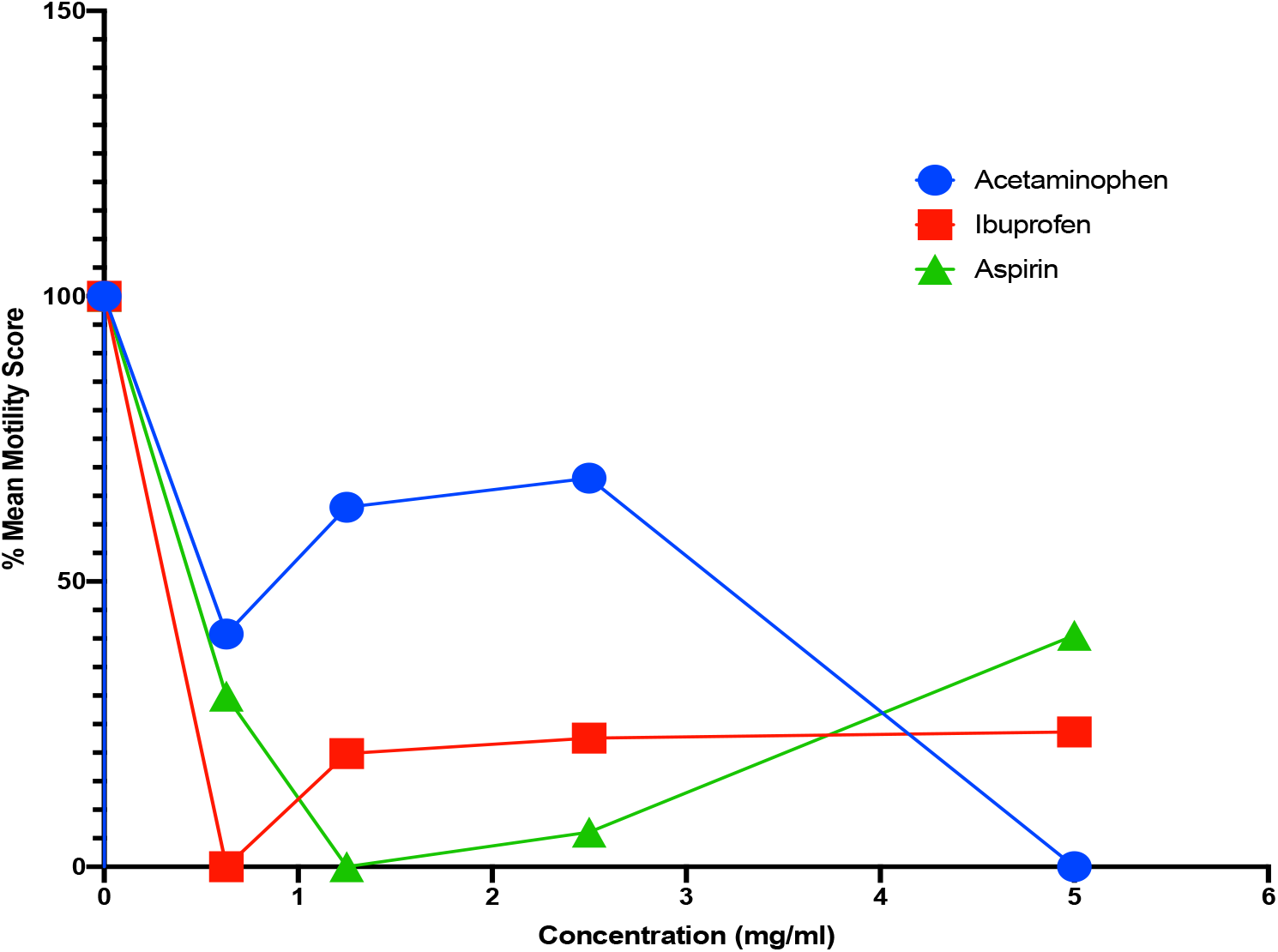
Dose response curve of the effects of Acetaminophen, Ibuprofen and Aspirin on female adult *O. volvulus* worms after 24 hours of incubation.

### Effects of acetaminophen, ibuprofen and aspirin on motility of adult worms

The results of the % MMS of the *O. volvulus* after a 24-hour exposure to the various concentrations of analgesics are presented in Figure 1. It can be seen in Figure 1 that at the 0.63 mg/ml dose of acetaminophen, the % MMS was 41% (SD=11.6%). The value of % MMS continued to increase as the concentration of acetaminophen increased [1.25 mg/ml (63%, SD=100%), then 2.5 mg/ml (68%, SD=18%)]. The only concentration of acetaminophen which performed best in terms of reducing motility of the worms was the 5 mg/ml, where the % MMS decreased to 5% (SD=0%). The EC50 (the concentration of a drug that gave half-maximal response) value of acetaminophen in this study was 8.74 mg/ml (57.82 mM).

The results show that at the lowest concentration of ibuprofen (0.63 mg/ml), the % MMS of worms in the ibuprofen group was 0% (SD=7%) compared to 100% (SD=100%) in the control group. For ibuprofen concentrations of 1.25 mg/ml, 2.5 mg/ml and 5 mg/ml, the % MMS of the female adult worms were respectively at 20 % (SD=0%), 23 % (SD=1%) and 24 % (SD=17%). The EC50 for ibuprofen was 0.65mg/ml (3.15 mM).

The findings on the effects of aspirin on the worms showed that at the startup aspirin concentration (0.63 mg/ml), 30 % (SD=61 %) of the worms were markedly motile within the aspirin group compared to 100 % in the control group. The lowest % MMS for worms in the aspirin group (0%, SD=0 %) was observed at an aspirin concentration of 1.25 mg/ml. However, at the 5 mg/ml concentration of aspirin, 41 % (SD=55 %) of the female adult worms in the group were still motile. The EC50 of aspirin in this study was 0.26 mg/ml (1.44mM).

### Response of O. volvulus parasites to varying concentrations of the test drugs during 8 days of culture

The following Figures (2–5) presents the results of the effects of varying concentrations of the test drugs (control, 5 mg, 2.5mg, 1.25mg/ml and 0.63 mg/ml) on female adult worms in culture for 8 days. Figure 2 summarizes the response of the of *O. volvulus* worms to 5 mg/ml concentrations of acetaminophen, ibuprofen, aspirin and control. The findings demonstrate that at 24 hours of culture (day 2), all test drugs including control had no effects on the motility of the worms as the % MMS of female adult worms in each group was 100 %. A similar observation was made on day 3 as all test drugs again had similar effects on worm motility. For day 4, the study observed that acetaminophen had the lowest effect on worm motility with % MMS of 21% (SD = 1.6%), with this trend continuing even to the 8^th^ day of culture where 55% (SD=5.7%) of worms were still highly motile despite exposure to 5mg/ml of acetaminophen. Generally, the differences in % MMS between the aspirin and control were statistically significant (P = 0.026; Mann-Whitney test). There was however no statistically significant difference in % MMS between acetaminophen and control (P = 0.052; Mann-Whitney test) and ibuprofen and control (P = 0.127; Mann-Whitney test).

**Figure 2:**
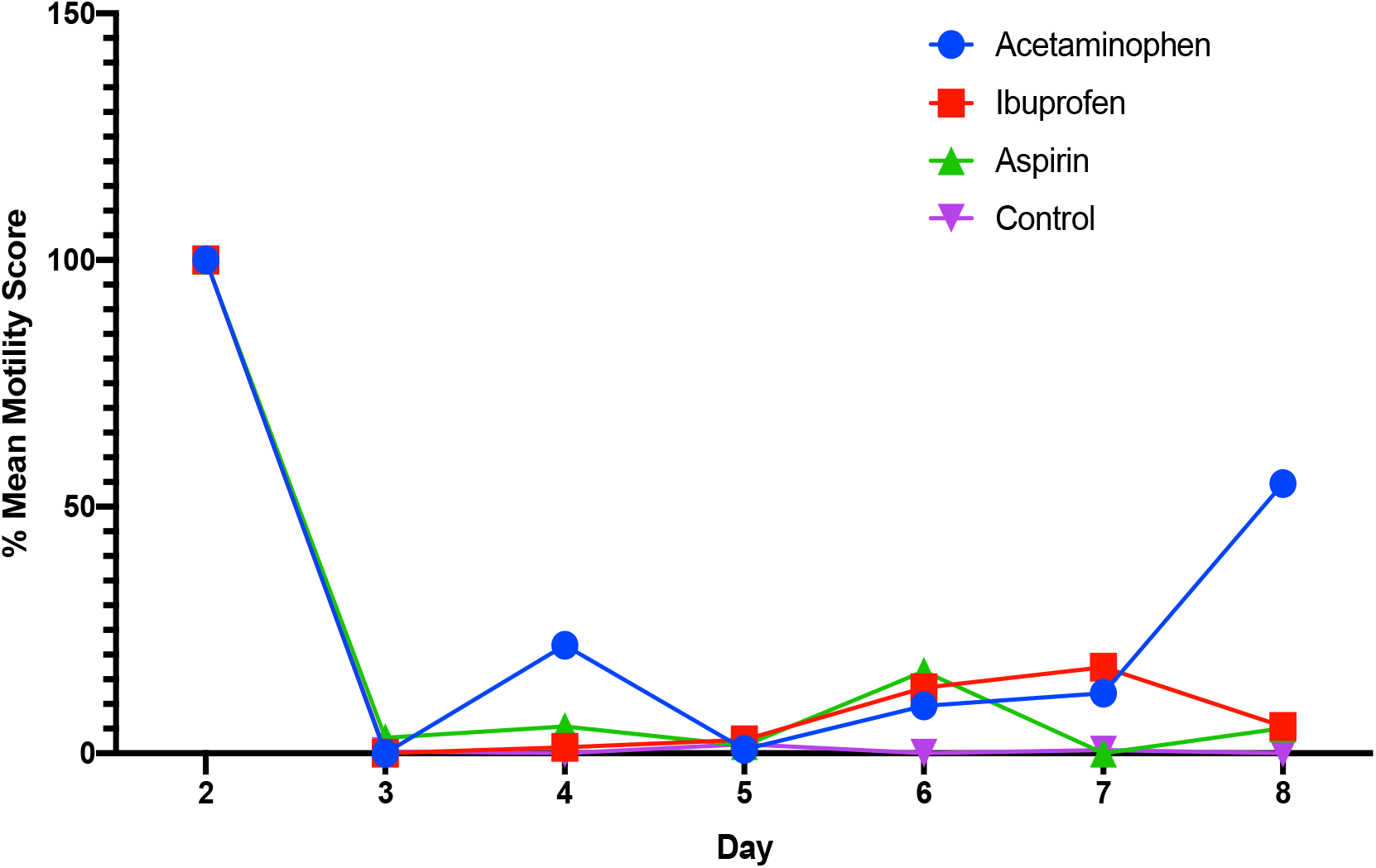
Comparison of 5 mg concentration of Acetaminophen, Ibuprofen, Aspirin on the motility of *O. volvulus* over the course of 8 days of culture.

A summary of the effects of the 2.5 mg/ml concentration of acetaminophen, ibuprofen and aspirin on the motility of female adult *O. volvulus* are presented in Figure 3. After 24 hours (day 2) of exposure of female adult *O. volvulus* to the test drugs, aspirin had the highest effect of reducing the % MMS, but even then, still had a % MMS of about 59% (SD = 2.2%). For the remaining days of culture (except on day 7), acetaminophen and ibuprofen outperformed aspirin. For instance, on day 3 of culture, the % MMS of the acetaminophen and ibuprofen group were only 5 % (SD = 56%) and 11% (SD = 16%) of worms in the, respectively, were motile compared to 30% of the worms in the aspirin group. It is interesting to note that on this day, the control group performed better than all the test drugs in terms of reduction in % MMS. There was a statistically significant difference between ibuprofen and control (P = 0.037; Mann-Whitney test) as well as between aspirin and control (P = 0.026; Mann-Whitney test). However, the difference between acetaminophen and control for this concentration group was not statistically significant (P = 0.104; Mann-Whitney test).

**Figure 3:**
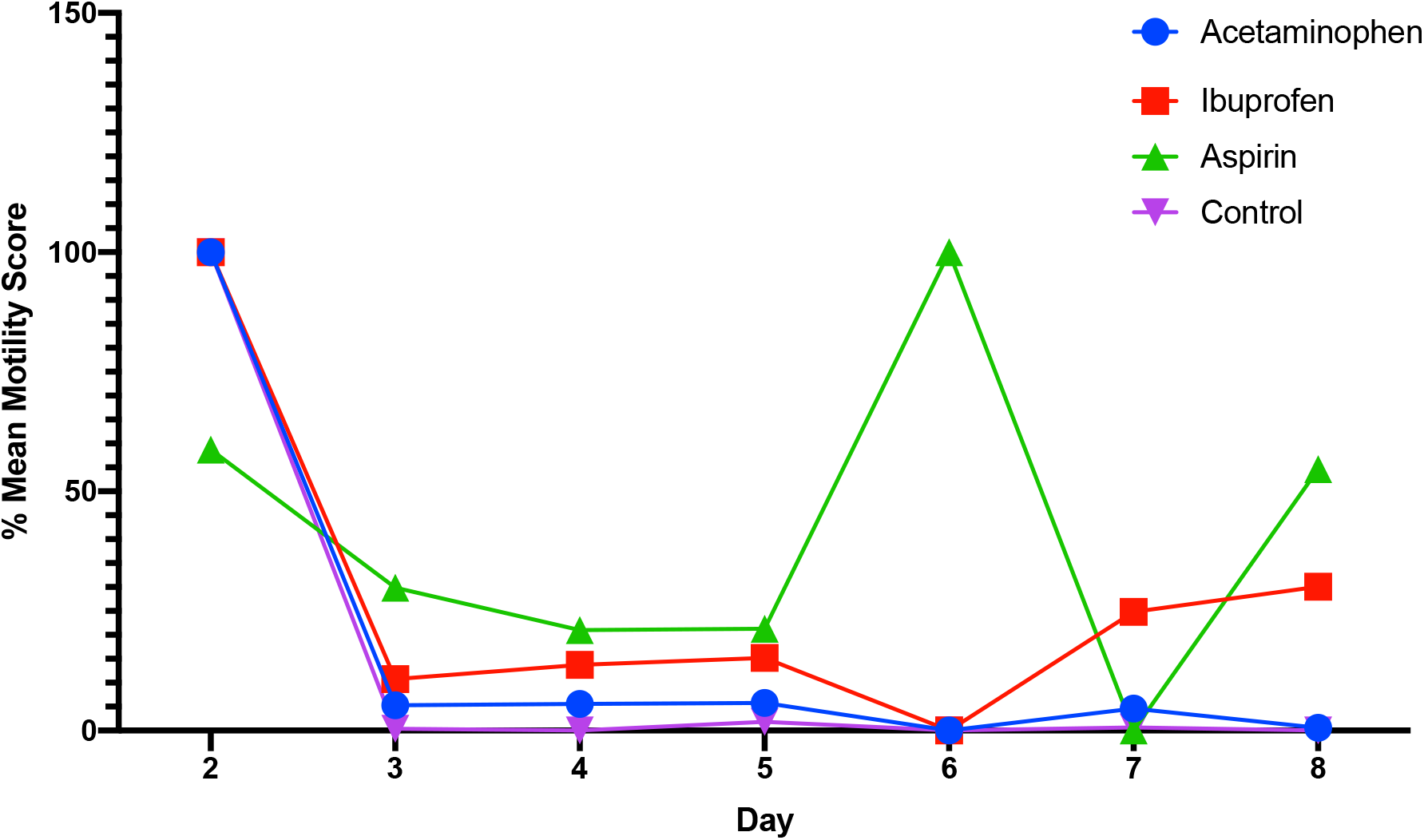
Comparison of 2.5mg concentration of Acetaminophen, Ibuprofen, Aspirin on the motility of *O. volvulus* over the course of 8 days of culture.

For the 1.25 mg /ml concentrations of the test drugs, aspirin performed best on day 2 (after 24 hours of exposure to the drug) though again, it was subsequently outperformed by all other test drugs including control (Figure 4). The only exception was on day 6, when no motility was observed for the aspirin group (0 % MMS, SD = 0 %) and day 8, when it performed better than ibuprofen. Surprisingly however, the control group continued to demonstrate very low motility throughout the culture period. For this drug concentration, there were statistically significant differences between acetaminophen and control (P = 0.026; Mann-Whitney test), ibuprofen and control (P = 0.037; Mann-Whitney test) but not between aspirin and control (P = 0.071; Mann-Whitney test).

**Figure 4:**
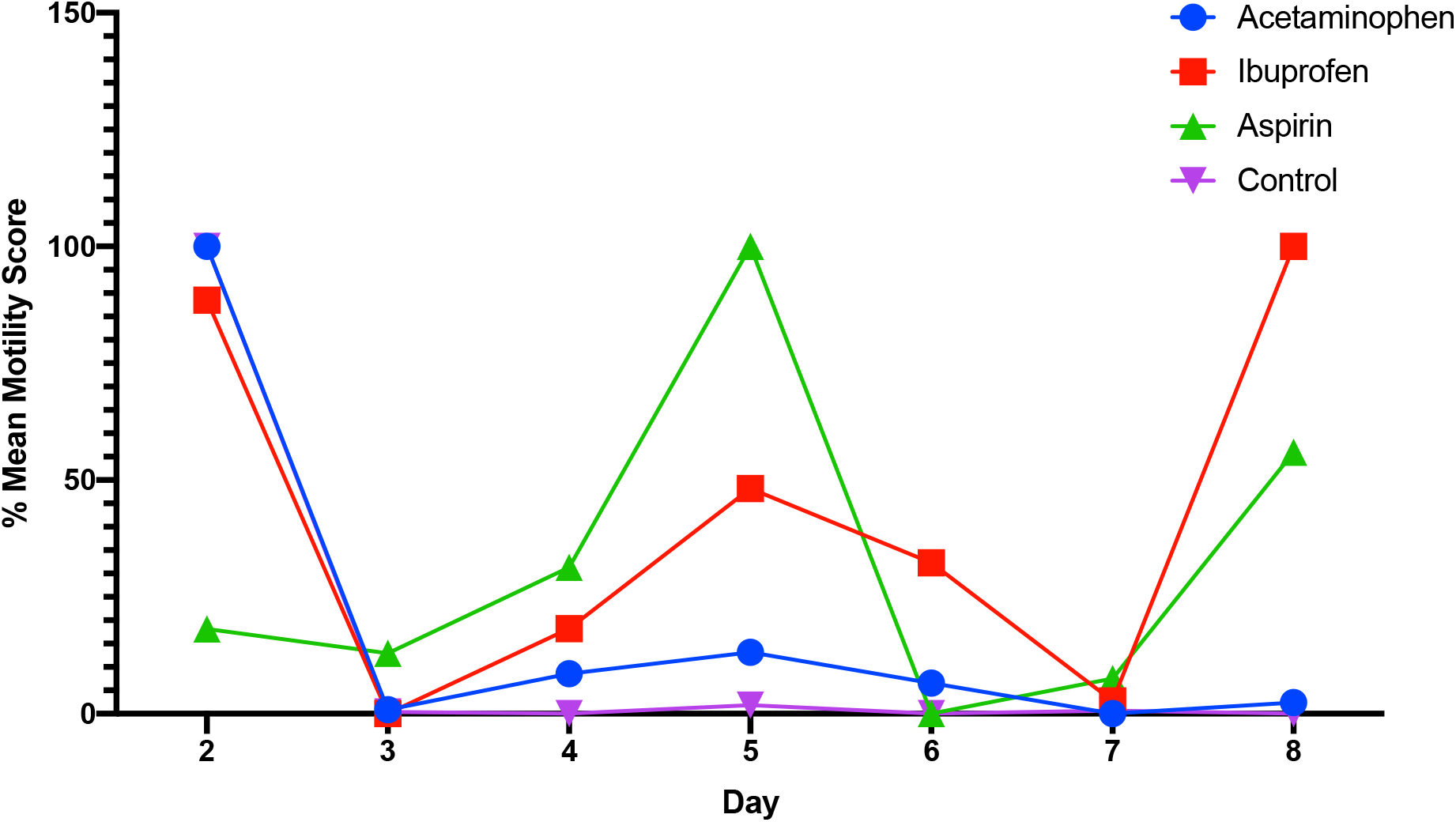
Comparison of 1.25 mg concentration of Acetaminophen, Ibuprofen, Aspirin on the motility of *O. volvulus* over the course of 8 days of culture.

The findings on the response of *O. volvulus* adult females to 0.63 mg/ml concentration of the various test agents as well as control is presented in Figure 5. On day 2, the best performing drug in terms of reduction in % MMS was ibuprofen, but even then, the % MMS was 70% (SD=7%) indicating a high worm motility in the group. Generally, however, acetaminophen performed better at this concentration when compared with aspirin and ibuprofen. Again, the control group continued to show relatively lower motility scores when compared with the acetaminophen, ibuprofen and aspirin. These differences in % MMS was statistically significant for ibuprofen versus control (P = 0.026; Mann-Whitney test) but not for aspirin versus control (P = 0.052; Mann-Whitney test) and acetaminophen versus control (P = 0.104; Mann-Whitney test).

**Figure 5:**
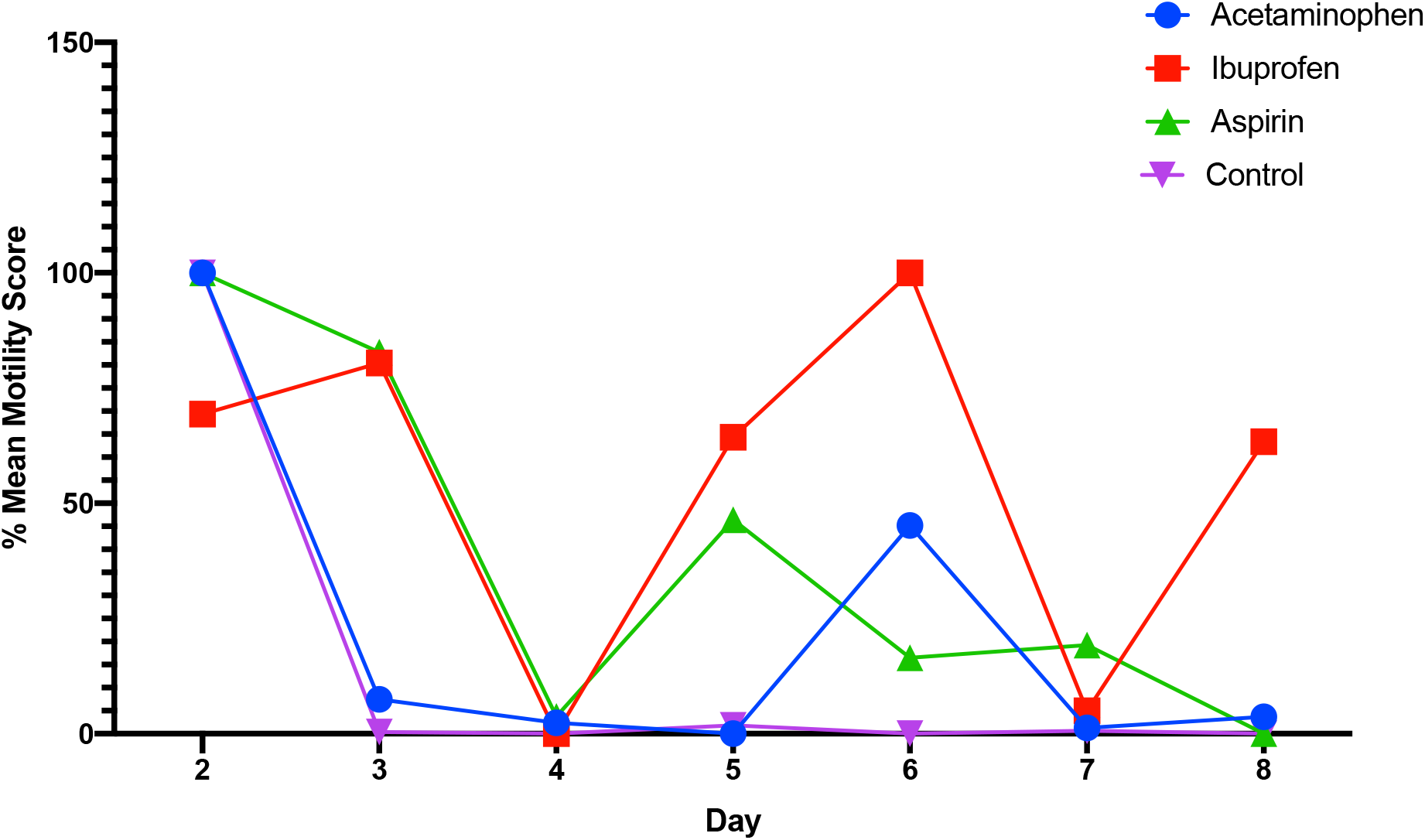
Comparison of 0.63 mg/ml concentration of Acetaminophen, Ibuprofen, Aspirin on the motility of *O. volvulus* over the course of 8 days of culture.

## Discussion

In response to the need to develop alternative drugs to augment ivermectin and based on empirical evidence from the sequencing of the genome of *O. volvulus* by Cotton et al.(2016), the present study tested the hypothesis that analgesics such as acetaminophen, ibuprofen and aspirin has adulticidal effects on adult female *O. volvulus* worms.

For acetaminophen, this study demonstrated that a the 24-hour exposure of the worms to 5 mg/ml acetaminophen completely immobilized the worms suggesting a potential anti-filarial effect. However, the summary of results of the various concentrations of acetaminophen over the course of the 8 days incubation revealed that all concentrations from this drug improved the motility of the female adult worms and hence did not indicate any anti-filarial effect. In fact, the 1.25 mg/ml concentration of acetaminophen significantly improved worm motility. This finding is contradictory to the suggestion by Cotton et al. (2016) that analgesics, such as acetaminophen, could have adulticidal effects on *O. volvulus* by blocking amidase, a potentially choke point enzyme. The ability of acetaminophen to enhance parasite motility and survival demonstrated in this study may not be surprising as a previous study on *Plasmodium falciparum* also demonstrated that the drug may prolong parasite clearance and had no direct antiparasitic effects [37]. Though the foregoing study was on malaria parasites, the results from the current study suggest a similar worm enhancing effect. It is therefore justified that prior to our study, acetaminophen had only been utilized as an antipyretic in the treatment of helminthic infections, with no direct effects on parasite intended. However, as the present study did not investigate the combination of acetaminophen and antifilarials, it is still possible that such a combination, in the right concentration, might be deleterious to *O. volvulus* parasite survival. This may not be far-fetched as some investigators have demonstrated the antimicrobial ability of acetaminophen on bacteria and fungi at different concentrations [23], [26]. Taken together, it is posited that there is a possible inhibitory effect of acetaminophen which is both dose and time dependent as the 24-hour exposure of worms to 5mg/ml acetaminophen completely immobilized them, but more research is needed in this area.

This study also investigated the effects of ibuprofen on the survivability of adult female *O. volvulus* worms. The findings showed that ibuprofen reduced the motility of the parasites at different concentrations within the first 24 hours of culture and this represented a two-fold increase in inhibitory activity over that of control. This finding is novel as this study represents the very first report of the inhibitory effects of ibuprofen on *O. volvulus* or on helminths in general. The antimicrobial effects of ibuprofen have been demonstrated previously for bacteria such as *Acinetobacter baumannii, Bacillus spp., Helicobactor pylori, Klebsiella spp*, etc as well as for fungi including *Candida albicans, Epidermophyton floccosum, Penicillium expansum* etc [26]. A rather surprising finding was the fact that summary of results of long-term (8 days) of exposure of worms to ibuprofen improved the motility of the worms. The improvement in worm motility by ibuprofen was statistically significant at all concentrations with the exception of the 5 mg/ml concentration when compared to the control group. This finding was particularly interesting because based on a previous study by Cotton et al.(2016), the current study had hypothesized that ibuprofen, being an analgesic, binds to amidase which is probably a chokepoint enzyme, leading to worm death. Thus, the reason for this observed significant improvement in motility of female adult worms following a relatively long-term exposure to ibuprofen is not known though the idea that ibuprofen may improve the longevity of microbes is not new [38]. He et al. (2014) discovered that the common drug ibuprofen increases the lifespan of yeast, worms and flies (specifically *Saccharomyces cerevisiae, Caenorhabditis elegans* and *Drosophila melanogaster* respectively), indicative of conserved longevity effects. They opined that in budding yeast, ibuprofen’s pro-longevity action is independent of its known anti-inflammatory role [38]. They also revealed that the critical function of ibuprofen in longevity is to inhibit the uptake of aromatic amino acids, by destabilizing the high-affinity tryptophan permease. Nevertheless, in consideration of the differences in effects of ibuprofen in the short-term and after extended exposure in this study, it is suggested that effect of ibuprofen just like acetaminophen is both dose and time dependent. The findings from the present study therefore begs for more research in this area to unravel further details on the effects of ibuprofen on *O. volvulus* worms and its potential role either as a treatment option or as a supplement in culturing worms in the lab.

The study further investigated the effects of aspirin on female adult *O. volvulus* worms. The findings showed that in the short-term, aspirin also demonstrated a 2-fold inhibition of the motility of the worms compared to the control group. This finding was not surprising as aspirin is a known nonsteroidal anti-inflammatory drug which can inhibit prostaglandin H synthase and also induces apoptosis [30]. In fact, Singh and Rathaur (2010) working with *Setaria cervi*, a bovine filarial parasite, demonstrated that exposure of the worms to a combination of diethylcarbamazine plus aspirin at a concentration of 100 mM irreversibly paralyzed adult worms as well as microfilariae within 2 hours of exposure. Evidence from their study also showed that the worms might have been killed due to mitochondrial mediated apoptosis [30]. In another study by Ochoa-Maganda et al. (2020), aspirin showed a direct anti-giardial activity and affected the adhesion and growth of trophozoites in a time-dose-dependent manner [18]. However, for the present study, what was surprising in the aspirin group was the observation that over the course of the entire period of incubation, aspirin seemed to enhance the motility of the female adult worms. The reason for this is not known though aspirin has also long been a subject of research on pro-longevity in humans, thereby warranting further research in the area.

As observed in this study, the general trend is that at the onset of incubation, acetaminophen, ibuprofen and aspirin, demonstrated some inhibitory effects on female adult worms. However, after an extended period of exposure, the drugs improved the motility of the worms indicating that the effects of these drugs are both dose and time dependent. It is speculated, that at the beginning of the introduction of the drugs, the analgesics might bind to amidase [21] and inhibit the worms within 24 hours duration of exposure. It is however possible, that amidase might not be a chokepoint enzyme. Hence, the worms could switch to alternative metabolic pathways [17] which does not require amidase and might thus increase their survivability *in vitro*, thereby accounting for the observed enhancement in worm motility scores in this study.

It was further observed that the control group which had only medium showed the worst motility score from day 3 onward. This was quite unexpected and the reasons are not immediately apparent. It is possible that the addition of feeder cells such as monkey kidney cells (LLXC-NK2-cells) in the assay would have significantly improved the survival of worms in the control [39]. However, Townson and colleagues (1986) in their work demonstrated the survival of *O. volvulus* worm *in vitro* for over 15 days even without feeder cells. More so, in the present study, the worms in the drug arms survived better even in the absence of feeder cells, hence the reasons for the low survival in the control group after day 3 cannot be due to the absence of these feeder cells alone. It is speculated that the additional filing materials in the drugs could contribute to the worm’s metabolism and thus improve worm motility compared to control. Hence, it is recommended that future investigations utilize pure research grade drug powder to eliminate possible confounders in the tablets. Nevertheless, the present study provides the foundation for further studies on the potential repurposing of these drugs for the treatment of onchocerciasis.

## Conclusions

The study provides, for the first time, evidence on the direct effects of acetaminophen, ibuprofen and aspirin on the motility of female adult *O. volvulus* worms. The study has demonstrated that a 24-hour exposure of these parasitic worms to the study analgesic drugs results in significant inhibition in worm motility compared with the control group. Longer duration of exposure of the worms to the different concentrations of the study drugs however led to a rather surprising enhancement in motility of the previously immobile worms. There is therefore the need for more research to determine the utility of these analgesics as possible anti-filarial drugs or supplements to improve *in vitro* culturing of *O. volvulus* worms for other studies.

## Data Availability

The datasets used and/or analyzed during the current study are available from the corresponding author on reasonable request.

## Acknowledgments

We thank all the staff and students of the Department of Basic and Applied Biology who provided support in one way or the other to ensure the successful implementation of this research. We also appreciate the efforts of the directors and staff of the Wenchi Municipal Health Directorate, Nsawkaw District and the Subinso Health Centre for their enormous contributions towards the study. Our appreciation also goes to the chiefs and people of the study communities who partnered with us in this study.

## Supplementary Materials

Not applicable

